# Transcription-dependent phase coexistence of mitochondrial nucleoids and RNA granules

**DOI:** 10.64898/2025.12.19.694874

**Authors:** Nidhi Parikh, Surya Teja Penna, Eliza Neal, Marina Feric

## Abstract

The spatiotemporal organization of multiple components within biomolecular condensates helps coordinate gene expression. Mitochondria separate transcription and RNA processing into two distinct condensates: mt-nucleoids and mtRNA granules (MRGs), respectively. However, how mtRNA transcripts are transferred from mt-nucleoids to distant MRGs was unclear. With high-resolution imaging, we examined the steady-state organization of mt-condensates in human cells. We identified a wide distribution of distances between centroids of mt-condensates, from roughly a micron apart to within 100 nm of each other. Live imaging revealed that such organization was dynamic: mt-condensates frequently underwent cycles of mixing and demixing. Indeed, mtRNA transcripts and the mtRNA polymerase co-localized within mixed mt-condensates, while transcription inhibition led to complete dissolution of MRGs, supporting that nascently transcribed mtRNA is a key driver of mt-condensate organization. Together, our results show that active transcription sustains the phase coexistence between mt-nucleoids and MRGs, with implications for transcriptional condensates more broadly.

## Introduction

The eukaryotic nuclear genome is hierarchically organized, allowing for spatiotemporal compartmentalization of gene expression^1–3^. Many assemblies of multiple transcriptional components appear as ∼100 nm sized puncta with liquid-like behavior, arising from intracellular phase transitions, and collectively, are referred to as transcriptional condensates^4–8^.

The emergent behavior–from their structure to their dynamics–of transcriptional condensates varies depending on the biological context^9^. For example, the tri-partite nucleolus is one of the most prominent condensates in the nucleus, where it nucleates around ribosomal DNA (rDNA) to synthesize ribosomal RNA (rRNA)^10^. RNA Polymerase I (Pol I) concentrates in the nucleolar fibrillar center and transcribes high levels of rRNA transcripts that get further processed in the dense fibrillar component before assembling into pre-ribosomal particles in the granule component^11^. Indeed, this layered, multi-phase organization arises from the degree of (im)miscibility between components from each sub-compartment and contributes to the outward flow of rRNA^12–15^

In contrast, all protein-coding RNA is synthesized by Pol II, associating with both dynamic and stable clusters that coordinate bursts of transcription throughout euchromatin^16^. These assemblies represent transcriptional condensates^17,18^ arising from transcription factors that assemble on promoter and enhancer sequences guided by their DNA-binding domains and recruit additional components, including Mediator, transcriptional co-activators, and Pol II, particularly through their intrinsically disordered domains^19–23^. Splicing occurs co-transcriptionally in the nucleus, and phosphorylation of the carboxy-terminal domain (CTD) of Pol II allows it to partition with splicing factors from associating nuclear speckles, effectively coordinating the transfer of RNA between transcriptional to splicing condensates^8,24,25^. For both Pol I and Pol II, transcriptional output appears as a key driver of the resulting phase behavior^9,26^; how nascent RNA contributes to the organization and stability of other transcriptional systems is still largely unknown.

In addition to the nuclear genome, eukaryotic cells also contain a second genome: the mitochondrial genome (mtDNA)^27^. The mitochondrial genome is small (16 kb) and organized within discrete nucleoprotein complexes called mt-nucleoids^28,29^. Mitochondrial gene expression encompasses similar processes as seen with Pol I and Pol II in the nucleus, albeit with markedly different organization. For example, mitochondria contain a single mitochondrial RNA polymerase (POLRMT) that transcribes both rRNA and mRNA as long, polycistronic transcripts^30–32^. The nascent mtRNA becomes processed in an entirely separate ribonucleoprotein complex – the mtRNA granule (MRG) – which are thought to be positioned a micron apart^33,34^. Indeed, mt-nucleoids^35,36^ and MRGs^37^ have recently been shown to exhibit features of biomolecular condensates. Thus, similar biological processes that occur in the nucleolus and in Pol II transcriptional condensates occur in the mitochondria, but each with unique apparent organization. How then does the coordinated transfer of mtRNA occur between these two seemingly disparate condensates within mitochondria? Here, we performed high-resolution microscopy to simultaneously visualize both mt-condensates in the context of transcription. We identified a transient sub-population of mt-nucleoids and MRGs that appeared well-mixed, and in this way, could directly coordinate the flow of mitochondrial gene expression.

## Results

### Endogenous mitochondrial condensates and steady-state mtRNA display a wide distribution of spatial localization

We first examined the endogenous spatial localization of mt-nucleoids, MRGs, and steady-state mtRNA transcripts simultaneously using high-resolution Airyscan imaging in fixed HeLa cells (Fig. 1a-e). Specifically, we labelled mtDNA and GRSF1 using immunofluorescence to mark mt-nucleoids and MRGs, respectively; and coupled with RNA FISH, we identified ND6 mt-mRNA transcripts. All components visualized appeared as distinct ∼100 nm foci distributed throughout the mitochondrial network (Fig. 1a), consistent with previous reports^37–41^.

**Fig 1.**
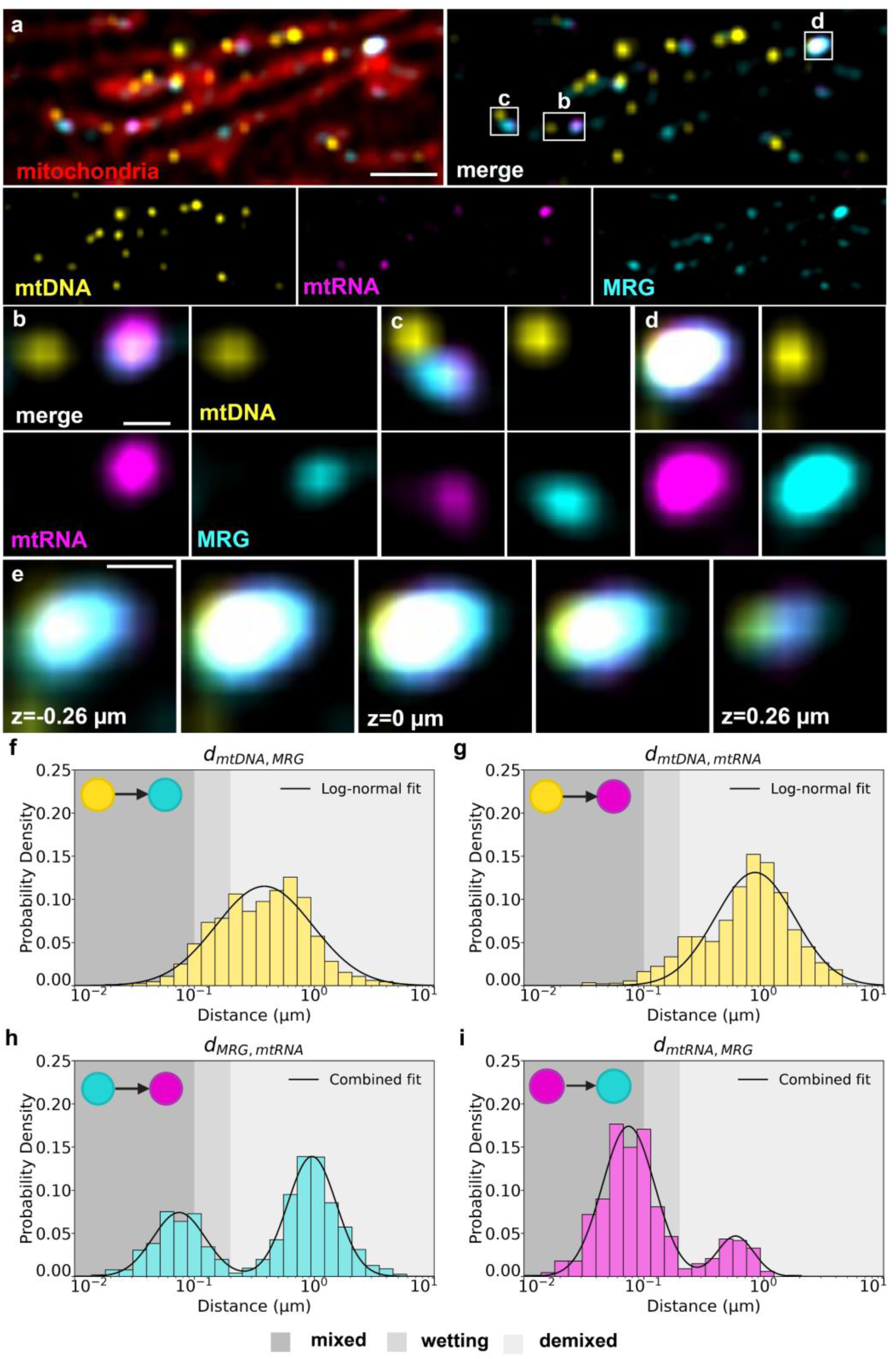
Endogenous localization of mitochondrial condensates and steady state mtRNA. **a**, Single *Z*-slice representative AiryScan image of a fixed, wildtype HeLa cell with labelled mitochondrial network (MitoTracker Red, red), mt-nucleoids (anti-DNA, yellow), ND6 mtRNA (RNA FISH, magenta) and MRGs (anti-GRSF1, cyan). Scale bar = 1 µm. **b-d,** Zoomed in images corresponding to ROIs from **(a)**: demixed **(b)**, wetting **(c)**, and mixed **(d)** mt-condensates. Scale bar = 200 nm. **e**, *Z*-stack of the fully mixed condensates in **(d)** spanning ∼0.5 µm. Scale bar = 200 nm. **f-i,** Quantitative analysis of nearest neighbor distances between pairs of mitochondrial components. Semi-log plots of probability density distributions of nearest-neighbor distances: mtDNA punctum paired to nearest MRG **(f)**; mtDNA punctum paired to nearest ND6 mtRNA **(g)**; MRG punctum paired to nearest ND6 mtRNA **(h)**; and ND6 mtRNA punctum paired to nearest MRG **(i)**. Shading of dark gray, medium gray and light gray indicate mixed, wetting and demixed populations, respectively, of the pairs of mitochondrial components within each plot (n=8 cells).

We next developed a quantitative image analysis pipeline to measure the nearest neighbor distances between puncta, *d_1,2_*, where *1* is the component of interest and *2* is its nearest neighbor, by identifying their centroids at sub-pixel resolution (see Methods). For example, *d_mtDNA,MRG_*, described the Euclidean three-dimensional distance of the nearest MRG to a given mt-nucleoid. We performed this analysis for each identified punctum for a given component and plotted the probability distributions (Fig. 1f-i, Extended Fig. 1c-d). We noticed several key features. First, there was a wide distribution in distances, suggesting their relative spacing is continuous. On average, mt-nucleoids and MRGs were ∼1 μm apart (*d̄*_*mtDNA,MRG*_= 0.54 +/- 0.01 μm SEM, n = 1,486 mt-nucleoids paired to nearest MRG); for many of these pairs, mtRNA was primarily concentrated in MRGs and not associating with mt-nucleoids (Fig. 1b,f). These results indicate a large majority of mt-nucleoids and MRGs were spatially separated (74%), where mtDNA was spaced more than 200 nm apart from mtRNA-containing MRGs (Fig. 1b,f).

However, there was a subset of the population that exhibited contrasting behavior. For example, we identified events where mtDNA and GRSF1 puncta, were spatially close: *d_mtDNA,MRG_* <200 nm (19%) (Fig. 1c,f). These events corresponded to neighboring mt-nucleoids and MRGs in our images (Fig. 1c). Lastly, we detected interactions at even smaller distances, of less than *d_mtDNA,MRG_* <100 nm (7%) and *d_mtDNA,mtRNA_* <100 nm (2%), suggesting complete overlap of these components (Fig. 1d,f,g). Consistently, these distances represented full-colocalization of all components as observed in our imaging (Fig. 1d). We confirmed this co-localization occurred in the same *Z*-plane by providing a montage of ∼0.5 um thick section (Fig. 1e, Extended Fig. 1a-b, Supplementary Video 1). Together, these findings indicate that although a majority of mtRNA-containing MRGs are largely demixed from mt-nucleoids, there are smaller populations that have significantly closer spatial associations. Such behavior is reminiscent of partially miscible liquid droplets that can undergo a range of wetting behavior^12,42^. We have thus characterized distances, *d* < 100 nm apart, as corresponding to well-mixed condensates, distances, 100 < *d* < 200 nm, as wetting, and distances greater than *d > 200* nm as demixed. Together, these results demonstrate a wide distribution in the relative spatial organization of mt-nucleoids and MRGs, spanning complete separation to full mixing.

### Spatially mixed mitochondrial condensates colocalize with active transcription

The range in spatial positioning of mitochondrial condensates relative to steady-state ND6 mt-mRNA transcripts was intriguing, and we hypothesized that the mixing events may be associated with active transcription. Thus, we next performed a series of experiments to identify active transcription. First, we investigated the colocalization of nascently transcribed and/or processed polycistronic mtRNA relative to mt-condensates by pulse labelling cells with modified nucleotide EU for one hour followed by immunofluorescence and click-chemistry based detection of EU (Fig. 2, Extended Fig. 2). We note here that the signal from EU represents total RNA, from both the heavy and light strands, including ribosomal and messenger mitochondrial RNA (rRNA, mRNA), in contrast to our previous measurements in which a specific transcript (ND6) from the light strand was labelled (Fig. 1). We observed largely similar localization patterns of EU-labelled mtRNA (Fig. 2a-d) to that of steady state mtRNA (Fig. 1), capturing mixing, wetting, and de-mixing behaviors. In the majority of cases, EU-labelled mtRNA was contained within MRGs, suggesting mtRNA is indeed quickly transferred to MRGs, consistent with previous reports^34,43,44^. However, we also saw similar wetting and mixing events with mitochondrial condensates, suggesting EU-labelled mtRNA also associated closely with mt-nucleoids (Fig. 2a, c, d). We quantified our observations using nearest neighbor distance analysis, where we detected a similar trend to the steady state mtRNA (Extended Fig. 2). Specifically, mt-nucleoids puncta were continuously distributed with average of ∼1 μm distances (*d̄*_*mtDNA,EU*_= 0.80 +/- 0.02 μm, SEM, n=1,112 mt-nucleoids paired to nearest EU) from EU puncta. To better understand how newly transcribed transcripts interacted with mt-nucleoids, we calculated the correlation coefficient that a third component has a similar interaction to the pair (Fig. 2e). We found significantly high correlation that EU-labelled mtRNA and an MRG both associate near a mt-nucleoid (ρ_mtDNA_ = 0.540, 95% CI: [0.040, 0.043]) compared to the less coordinated behavior of ND6-labelled mtRNA and MRGs (ρ_mtDNA_ = 0.037, 95% CI [0.0510, 0.0512]) (Fig. 2e). This high correlation of EU (nascent) transcripts and MRG simultaneously both associating with mt-nucleoids supports the notion that transcription occurs during mixing (Fig. 2e). Collectively, these results suggest that the spatial coupling of the mitochondrial condensates is coordinated with active transcription.

**Fig 2.**
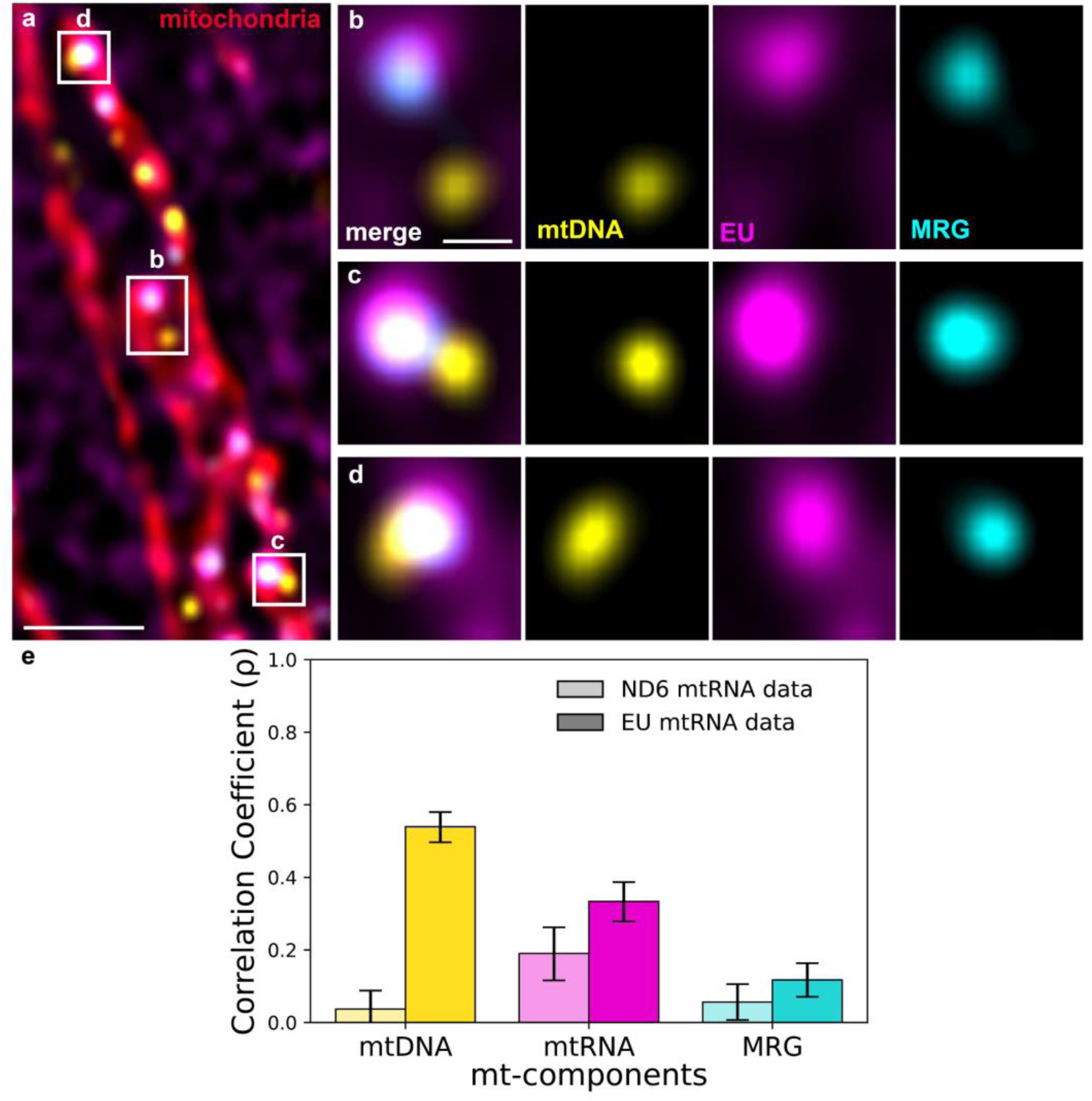
Colocalization of coexisting mitochondrial condensates and EU-labelled mtRNA. a, Single *Z*-slice representative AiryScan image of a fixed wildtype HeLa cell with labelled mitochondrial network (MitoTracker Red, red), mt-nucleoids (anti-DNA, yellow), nascent RNA transcripts (EU, magenta) and MRGs (anti-GRSF1, cyan) (n=10 cells). White box indicates four channel overlays of ROI. Scale bar = 1 μm. **b-d,** Zoomed in images ROIs of demixed (**b**), wetting (**c**), and fully (**d**) mixed mt-condensates in the context of EU. Scale bar = 200 nm. **e,** Correlation coefficients representing the likelihood of two mitochondrial components having similar nearest neighbor distances relative to the indicated component. Lighter colors show correlation coefficients for the ND6-labelled mtRNA (steady-state) data from Fig. 1, while darker colors show correlation coefficients for the EU-labelled mtRNA data (nascent). Error bars denote 95% confidence intervals. P-values comparing steady-state and nascent correlation coefficients are as follows: p=6.6E-46 for n=1,466 mtDNA puncta (ND6) and n=1,111 mtDNA puncta (EU); p= 0.0019 for n=668 mtRNA puncta (ND6) and n=1,035 mtRNA puncta (EU); p= 0.078 for n=1,568 MRG puncta (ND6) and n=1,737 MRG puncta (EU).

### Mitochondrial RNA polymerase partitions with both mitochondrial condensates

To further resolve if active transcription is occurring upon mixing, we sought to visualize the mitochondrial RNA polymerase, POLRMT. We generated an overexpression construct of POLRMT fused with HaloTag to visualize its spatial localization within the mitochondrial network. Upon 48 hours of POLRMT-HaloTag overexpression, we subsequently labelled the mt-condensates and ND6 mt-mRNA. We noted that overexpression of POLRMT did not appear as punctate as other markers of mitochondrial condensates as there was significant signal throughout the mitochondrial matrix. Nevertheless, POLRMT appeared to co-localize with both mt-condensates (Fig. 3a, Extended Fig. 3). As POLRMT-HaloTag did not form discrete puncta, we were unable to apply our nearest-neighbor analysis for accurate centroid detection; instead, we quantified the degree of partitioning by performing averaging of POLRMT signal centered on specific mitochondrial components^18^. For example, we identified every mt-nucleoid punctum and averaged the signal of all channels to produce a representative heatmap (Fig. 3b). Quantification of the average line profile from our heatmaps revealed that, on average, mt-nucleoids were partially enriched in POLRMT with low partitioning of MRGs and mtRNA (Fig. 3c). This analysis applied on mtRNA showed inverse behavior, where MRGs associated more strongly mtRNA with partial enrichment of POLRMT and weak localization of mtDNA (Fig. 3d,e). Indeed, these results are qualitatively consistent with the different populations identified from our nearest-neighbor analysis (Fig. 1). These findings indicate POLRMT partitions in both mt-nucleoids and MRGs, potentially functionally coupling the mitochondrial condensates when mixed.

**Fig 3.**
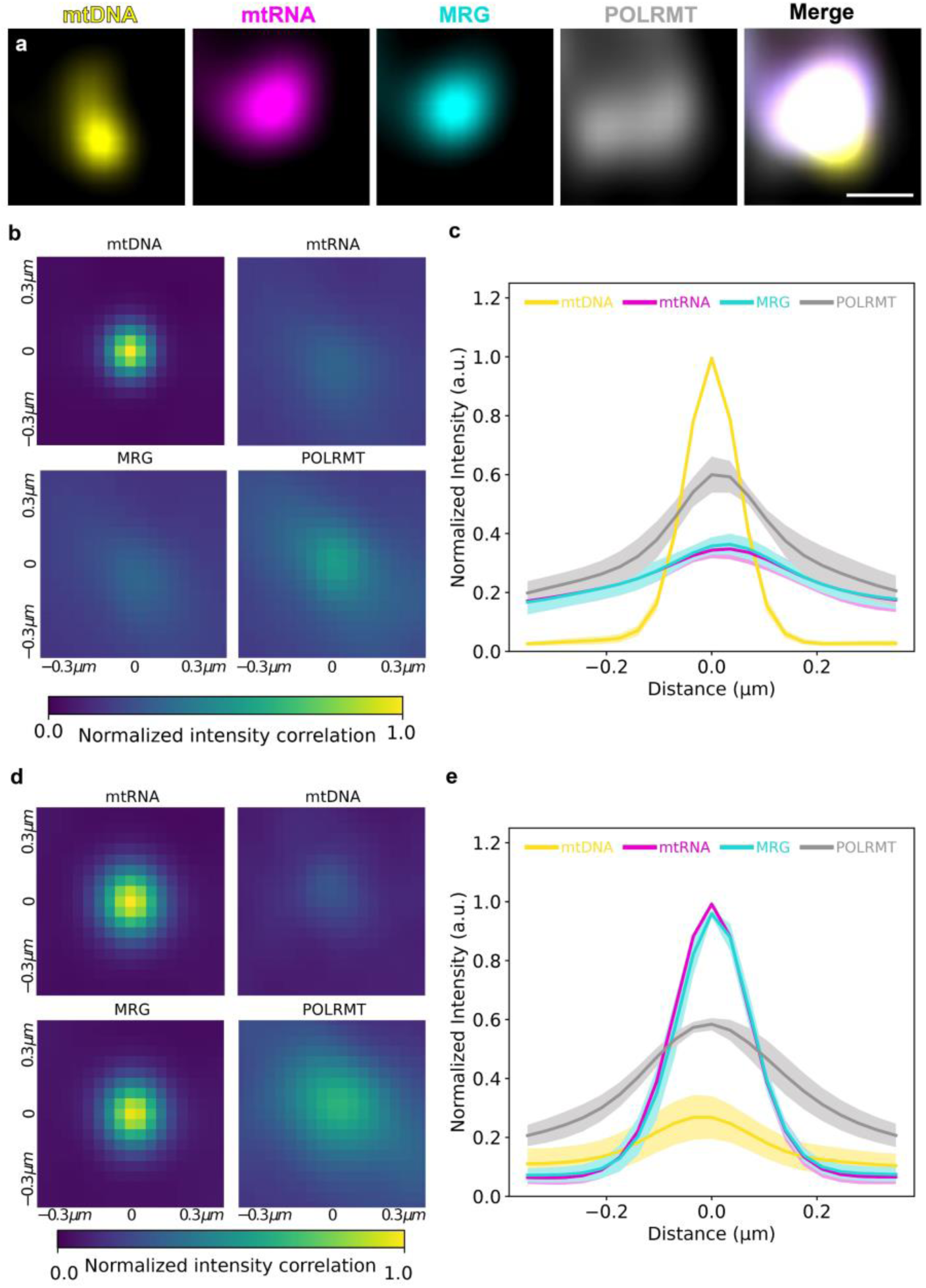
Colocalization of coexisting mitochondrial condensates and POLRMT. **a**, Single *Z*-slice representative AiryScan images of co-localizing mitochondrial condensates in a fixed wildtype HeLa cell over-expressing POLRMT-HaloTag (Oregon-Green-ligand, gray) with labelled mt-nucleoids (anti-DNA, yellow), ND6 mt-mRNA (RNA FISH, magenta), and MRGs (anti-GRSF1, cyan). Single channel of mtDNA, mtRNA, MRG, and POLRMT followed by four-channel merge. Scale bar = 200 nm. **b,d**, Average intensity of all channels based on identified mtDNA **(b)** and mtRNA **(d)** puncta in one cell. **c,e,** Line profiles across the centerline for all identified mtDNA **(c)** and mtRNA **(e)** puncta (error bar = s.d., n=10 cells).

### Mitochondrial condensates undergo dynamic mixing events concomitant with POLRMT partitioning

To determine whether mt-nucleoids and MRGs persist as distinct populations (i.e. mixed, wetting, or demixed) or rather dynamically sample multiple organizations, we performed multi-channel time-lapse imaging of live cells (Fig. 4). We visualized mt-condensates by labelling mt-nucleoids with dsDNA-intercalating dye PicoGreen^35^ and by overexpressing FASTKD2 fused with mScarlet to label MRGs^44^. Additionally, we overexpressed PORLMT fused to HaloTag (POLRMT-HaloTag) as a representative marker of mitochondrial transcription. Within a few minutes, we captured multiple cycles of rapid mixing, wetting and de-mixing of the two mt-condensates (Fig. 4a-e, Extended Fig. 4, Supplementary Video 2). We quantified these events by tracking the centroids of representative pairs of mt-condensates, and we plotted their 2D Euclidean distances over time (Fig. 4f). For example, one pair remained well mixed (<100 nm) ∼15% of the imaging time by undergoing transient interactions. Moreover, as POLRMT tended to be more diffuse in the mitochondrial network (Fig. 3), we frequently observed strong partitioning with both mt-nucleoids and MRGs, particularly during mixing events (Fig. 4b,e). Together, these findings further support the mechanism that mixing events are linked to active transcription.

**Fig 4.**
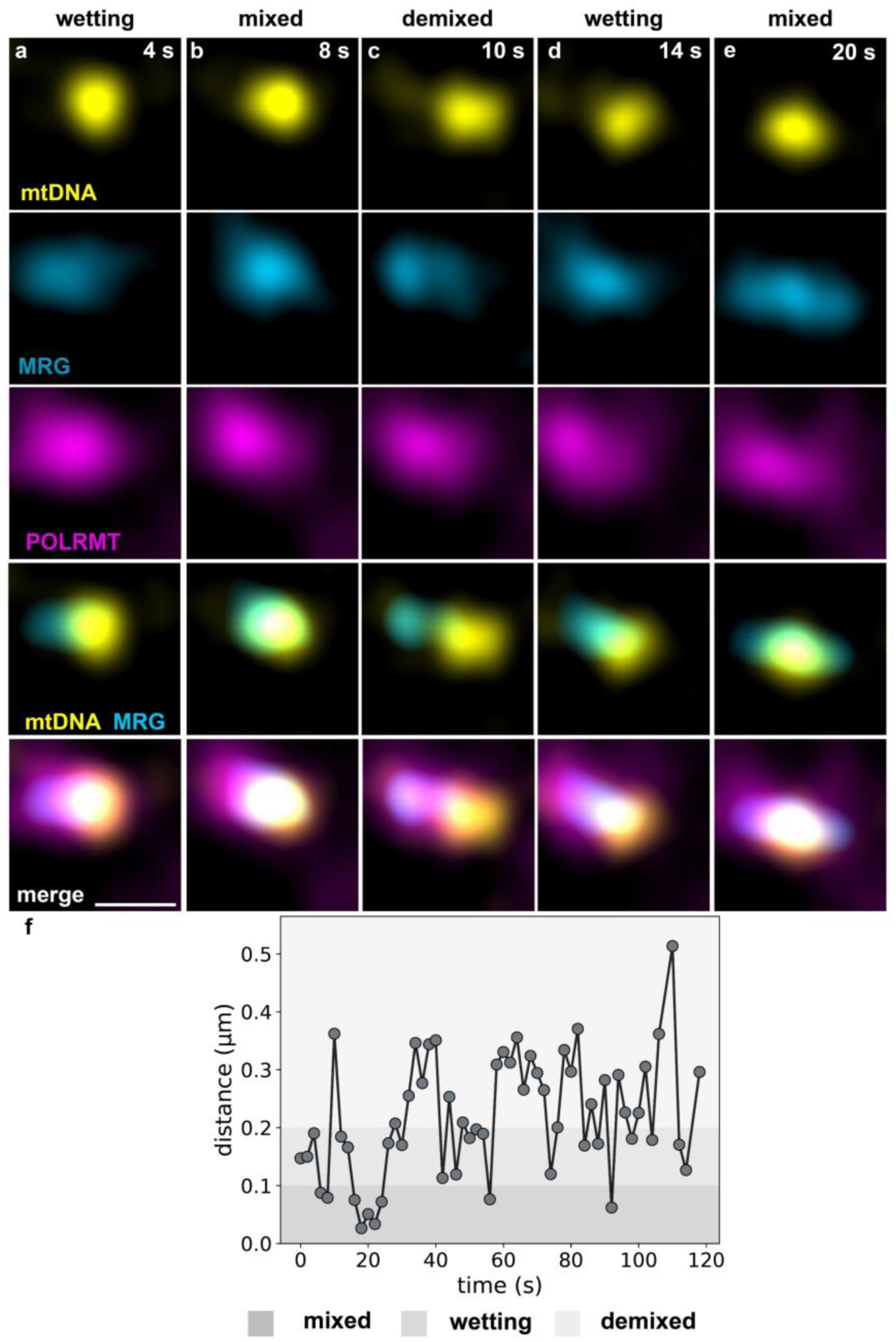
Dynamic mixing and de-mixing of mitochondrial condensates in live cells. **a-e**, Single *Z*-slice, time-lapse Airyscan imaging of a live wildtype HeLa cell labelled with mt-nucleoids (PicoGreen, yellow), mt-RNA granules (FASTKD2-mScarlet, light blue), and mtRNA polymerase (POLRMT-HaloTag, magenta). Scale bar = 500 nm. **f**, Quantification of change in 2D Euclidean distance between centroids of a representative pair of mt-condensates (mt-nucleoid and MRG) with time (n=14 cells).

### Small molecule inhibition of POLRMT leads to dissolution of MRGs

To directly assess the role of mitochondrial transcription in contributing to the spatiotemporal distribution of mitochondrial condensates, we treated cells with IMT1B, the specific small-molecule inhibitor of POLRMT^45^ (Fig. 5, Extended Fig. 5). Upon six hours of IMT1B treatment, we found decreased intensity of ND6 mt-RNA puncta by high-resolution microscopy (Fig. 5a,b), consistent with the known inhibitory activity of IMTIB^45^. Strikingly, we also noted that GRSF1 appeared diffuse throughout the mitochondrial network, without its canonical punctate organization (Fig. 5a, b). Quantification of intensities using line profiles confirmed a reduction for both mtRNA and GRSF1, while mtDNA remained largely unaffected (Fig. 5c,d). These results indicate that reduced RNA levels–resulting from transcription inhibition–lead to the dissolution of MRGs without disrupting the stability of mt-nucleoids.

**Fig 5.**
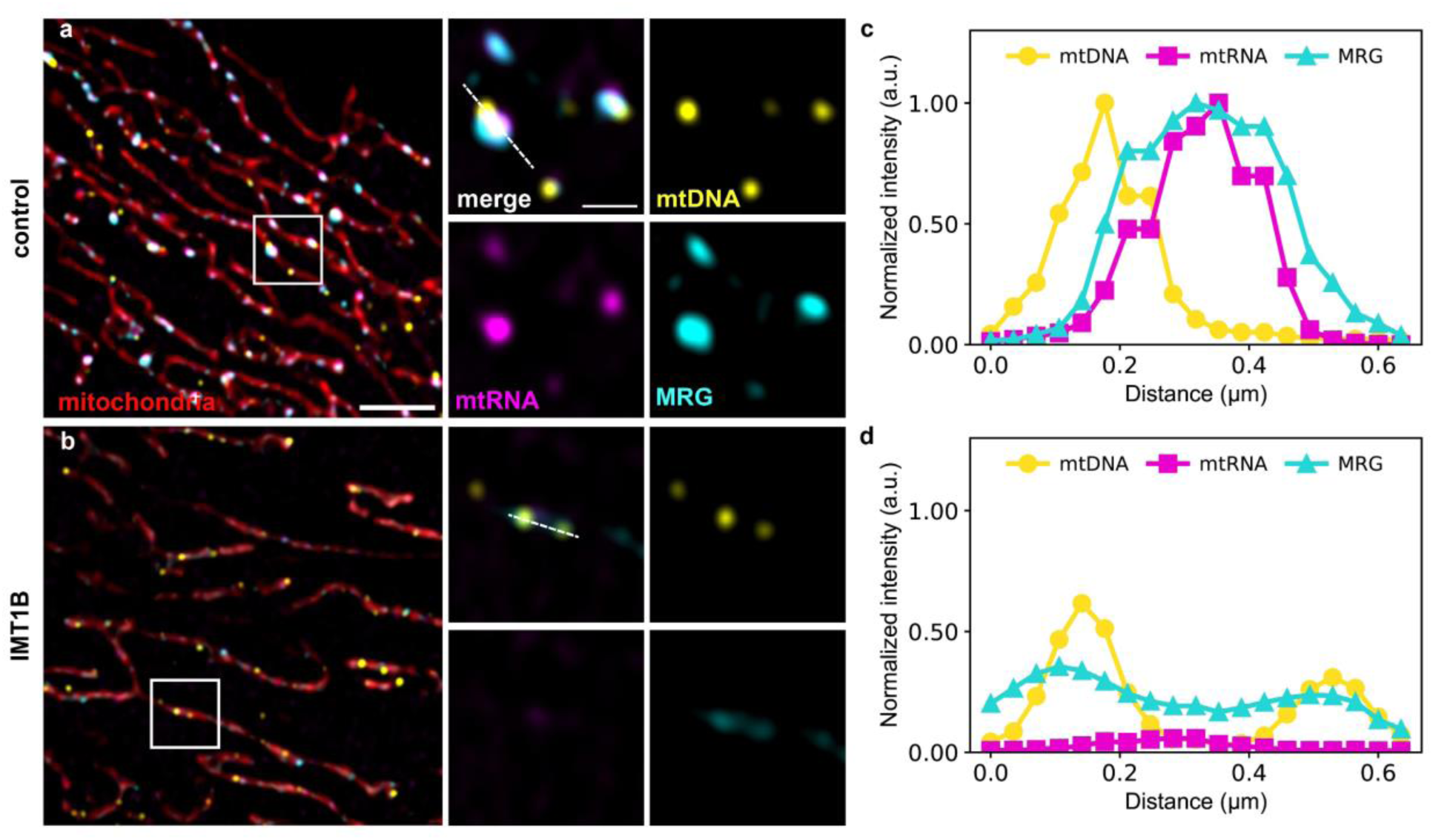
Dissolution of MRGs upon inhibition of mitochondrial transcription. **a,b**, Single *Z*-slice representative Airyscan images of a fixed wildtype HeLa cell treated with DMSO (negative control) **(a)** or 10 µM of IMT1B **(b)** for 6 hours with labelled mitochondrial network (MitoTracker Red, red), mt-nucleoids (anti-DNA, yellow), ND6 mt-mRNA (RNA FISH, magenta) and MRGs (anti-GRSF1, cyan). Scale bar = 2 μm. White boxes indicate regions of interest. Zoomed in images include a three-channel overlay and single channels of mtDNA, mtRNA and MRGs. Scale bar = 500 nm. **c,d,** Line profiles indicate normalized intensity distributions of mtDNA, mtRNA, and MRG corresponding to dashed lines on zoomed in ROIs from **a** and **b**, respectively (n=10 cells for control, n=11 cells for IMT1B treatment).

## Discussion

Our findings reveal that mitochondrial condensates – mitochondrial nucleoids and RNA granules – are highly dynamic structures that sample a range of spatial organizations as a function of transcriptional activity. We show that mt-condensates exhibit demixed, wetting and mixing behaviors, characteristic of multiphase droplets^46^. The close spatial interactions we observe are coordinated with transcription, explaining how RNA can be directly transferred from mt-nucleoids to MRGs for subsequent processing.

One theme that emerges is that the steady-state organization of mt-nucleoids and MRGs appears to be inherently dependent on transcription. Only a small fraction (∼15%) of mitochondrial nucleoids is reported to be transcriptionally active at any given time in human fibroblasts^47^. Consistently, we observe that a small fraction (∼9%) of mt-nucleoids that co-localize (<200 nm) with EU-labelled transcripts *and* MRGs, thereby linking close spatial proximity of the two condensates to transcriptional activity. Moreover, these events appear to be short-lived, as the mixed condensates rapidly separate, undergoing “kissing” events reminiscent of Pol II transcriptional condensates^17^.

We visualized a high enrichment of POLRMT within MRGs, which is intriguing given their primary site of activity is in mt-nucleoids. Indeed, immunoprecipitation techniques have also identified POLRMT in the proteome of mt-nucleoids^48^ as well as MRGs^33^, further supporting its association with both structures. Similarly, MRGs also concentrate two other transcriptional components, including transcription factor TFB1M^33,49,50^ and repressor MTERFD1^51–54^. These results point to MRGs also serving as a reservoir of key transcriptional components, and by mixing with mt-nucleoids, can regulate transcription in this way. Similar behavior is observed in other contexts; for example, transcriptional condensates provide components, including Mediator and transcription factors, needed to recruit and activate Pol II^18^ .

We showed here that the pharmacological inhibition of POLRMT disrupts MRG assembly, highlighting a crucial architectural role of RNA in MRG condensate formation and stability. RNA is well known to strongly influence the phase behavior in other transcriptional contexts^55–57^. RNA can undergo significant post-transcriptional modifications and can take on a variety of secondary structures, influencing its interactions with other RNA molecules as well as the associating proteins, and in this way, can shape the emergent behavior of condensates. Most strikingly, blocking rRNA production by inhibiting Pol I transcription, leads to large-scale rearrangements of nucleolar layers, suggesting continuous RNA production and processing are needed to maintain proper nucleolar architecture^14,15,58^. The most direct observations of the diversity in phase behavior arising from RNA are made through *in vitro* transcriptional systems. For example, *in vitro* reconstitution describe an RNA-dependent condensation or dissolution of Pol II^60^, formation of vesicle-like structures where DNA and RNA partitioned in different layers upon mitochondrial transcription^36^, or how orthogonal RNAs from a purely synthetic system can form co-transcriptionally into entirely immiscible droplets^61^. The network of interactions between mtDNA, mtRNA, and associated mt-proteins likely underlies the molecular mechanisms determining mt-nucleoid and MRG phase coexistence.

Our results together thus provide a model explaining how the phase behavior of mt-condensates contributes to the flow of mitochondrial gene expression. Steady-state mt-nucleoids and MRGs largely coexist as demixed condensates. Yet, they can diffuse within the mitochondrial matrix, and undergo wetting, during which the MRGs likely contribute key factors–particularly POLRMT–necessary to initiate transcription. Mixing further allows for exchange of key components, but as RNA transcripts are produced, the resulting RNA decreases the solubility of the condensates^36^, driving them apart. Thus, our observed sub-population of mixed mt-condensates likely represents the actively transcribing mt-nucleoids, primely positioned to rapidly hand off the nascent polycistronic transcript to the MRGs to be stabilized and further processed. Our results suggest nascently transcribed mtRNA serves as a key architectural molecule that regulates the spatial organization of mitochondrial condensates.

## Acknowledgements

We thank all members of Feric laboratory and PSU Center for Eukaryotic Gene Regulation (CEGR) for their feedback and discussion. We thank the Genomics Core Facility at the Huck Institutes of The Life Sciences at PSU for their whole genome-sequencing technologies (RRID: SCR_023645). Research reported in this study was supported by the National Institutes of General Medical Sciences of the National Institutes of Health under award R35 GM154931 (MF).

## Statistics and Reproducibility

Each experiment had at least three technical replicates (n ≥ 3 cells). The exact sample sizes in each experiment are reported in the figure legends. Each experiment in the main figures was repeated independently three times on different days. For comparison of nearest neighbor distances between the steady-state (ND6) and nascent (EU) data, distances were log-transformed before analysis. Two-tailed Welch’s t-test was applied to unimodal distributions, while two-tailed Mann-Whitney U test was applied to bimodal distributions. Pearson correlation coefficients (ρ) between mitochondrial components were calculated for both steady-state and nascent mtRNA datasets and compared using Fisher’s *r-to-z* transformation for independent samples. Two-tailed *Z*-tests were performed for each component, and 95% confidence intervals were calculated in Fisher z-space and back transformed to r. P-values were calculated with appropriate tests for each distribution and indicated in the figure legends.

## Data availability

Source data used to generate figures are provided with this paper and available through GitHub (https://github.com/fericlab/mt-condensates). All data supporting the findings of this study are available from the corresponding author upon reasonable request.

## Code availability

All the analysis scripts used in this study are available through the GitHub webpage at: (https://github.com/fericlab/mt-condensates).

## Contributions

M.F. and N.P. designed the study, discussed results, and wrote the paper. N.P. performed all high-resolution microscopy experiments and analyzed data. S.T.P. wrote code. E.N. contributed to IMT1B experiments. All authors reviewed the manuscript.

## Methods Cell

### Culture

HeLa cells (ATCC, CCL-2, Lot #70046455) were cultured in Dulbecco’s Modified Eagle Medium (DMEM, Thermo Fisher Scientific, #11960044) supplemented with 10% fetal bovine serum (FBS, Thermo Fisher Scientific, #A5256701) and 1% penicillin-streptomycin-glutamine (Thermo Fisher Scientific, #10378016) at 37°C with 5% CO_2_.

### Plasmid design and cloning

To generate FASTKD2-mScarlet, cDNA was synthesized from previously extracted RNA (human wild-type fibroblast HGMDFN090, Progeria Research Foundation)^1^ using RevertAid Reverse Transcriptase (Thermo Fisher Scientific, #EP0441). The FASTKD2 sequence was PCR-amplified from the cDNA, incorporating a C-terminal GGGGSx3 linker. The mScarlet sequence was amplified from pSL1088 plasmid (a gift from Scott Lindner) and positioned downstream of the linker. All fragments were assembled into the pmCherryN1 vector backbone (Takara Bio, #632523) using the NEBuilder HiFi DNA Assembly Cloning Kit. To generate POLRMT-HaloTag, POLRMT was PCR-amplified from cDNA (see above) with a C-terminal GGGGSx3 linker and cloned into the HaloN1 vector backbone (Halo-TOMM20-N-10, a gift from Kevin McGowan, Addgene #123284) such that HaloTag was positioned downstream of the linker.

### Immunofluorescence

Cells were seeded on coverslips in 12-well plates on Day 0. Immediately prior to fixation on Day 1, cells were treated with 100 nM MitoTracker Red CMXRos (Thermo Fisher Scientific) for ∼20 min at 37°C and 5% CO_2_ to label the mitochondrial network. Cells were fixed with 4% paraformaldehyde (PFA, Electron Microscopy Sciences) by diluting 16% PFA to 8% in 1X Phosphate Buffered Saline (PBS) and further directly diluting 1:2 into cell culture media for 20 minutes at room temperature, followed by rinsing with PBS. Cells were permeabilized with 0.5% Triton X-100 for 10 minutes, washed three times with PBS, and incubated with primary antibodies in PBS without blocking agents for 1 hour at room temperature. Primary antibodies included: anti-GRSF1 (1:1000, Sigma, HPA036985) and anti-DNA (1:500, Sigma, CBL186). Cells were washed three times with PBS, incubated with secondary antibodies for 1 hour at room temperature in the dark, and washed again three times with PBS. Secondary antibodies included: goat anti-rabbit Alexa Fluor 488 (1:1000, Thermo Fisher Scientific), goat anti-rabbit STAR Orange (1:500, abberior) and goat anti-mouse Alexa Fluor 405 (1:50, Thermo Fisher Scientific). Cells were post-fixed with 4% PFA for 10 min, rinsed twice with PBS. RNA labelling (EU or RNA FISH, see below) was subsequently performed before mounting onto slides with ProLong Gold Antifade Mountant (Thermo Fisher Scientific). Samples were cured for ≥48 hours at room temperature in the dark before imaging. Slides were stored at 4°C long-term. Additionally, for POLRMT-HaloTag localization, cells were transfected on Day 1 after seeding using FuGene HD Transfection Reagent (Promega, #E2311) following the manufacturer’s optimized instructions. On Day 3, cells were labelled with 0.5 μM HaloTag Oregon Green ligand (Promega, #G2802) for 1 hour in imaging media (DMEM without phenol red, Thermo Fisher Scientific #31053028), rinsed twice with media, and fixed before immunofluorescence and RNA FISH labelling.

### RNA FISH

Custom Stellaris RNA FISH probes targeting mt-ND6 were designed using the Stellaris RNA FISH Probe Designer (Biosearch Technologies, Petaluma, CA). Cells were hybridized with the Stellaris RNA FISH Probe set labelled with Quasar 670 (Biosearch Technologies) following manufacturer’s protocol for sequential immunofluorescence and RNA FISH. Briefly, fixed and permeabilized cells underwent immunofluorescence without blocking agents, were post-fixed with 4% PFA for 10 min, and hybridized with RNA FISH probes.

### EU incorporation

Cells were seeded on coverslips in 12-well plates on Day 0. On Day 1, cells were incubated with 2.5 mM EU (Thermo Fisher Scientific, #E10345) for 1 hour in DMEM. Following EU incubation, EU-containing media was removed, and cells were labelled with 100 nM MitoTracker Red for 30 min, immediately fixed with 4% PFA for 20 minutes, and processed for immunofluorescence followed by post-fixation. After post-fixation, cells were permeabilized again with 0.5% Triton X-100 and labelled for EU detection with 2.5 mM Alexa Fluor 647 Azide (Thermo Fisher Scientific, #A10277) using the Click-iT Cell Reaction Kit (Thermo Fisher Scientific, #C10269) following the manufacturer’s instructions, excluding blocking agents.

### IMT1B treatment

Inhibitor of mitochondrial transcription 1B (IMT1B) (Thermo Fisher Scientific, #HY-137067) was reconstituted in DMSO in accordance with manufacturer’s instructions. Cells were seeded on coverslips in 12-well plates 24 hours prior to treatment. Media was removed and replaced with untreated media, media containing DMSO (control), or media containing 10 μM IMT1B for six hours^2^. Cells were subsequently incubated with 100 nM MitoTracker Red for 30 minutes prior to fixation. Briefly, subsequent immunofluorescence with anti-GRSF1 and anti-DNA staining followed by RNA-FISH (see above) was performed.

### Transient transfection and live imaging

Cells were seeded in 8-well imaging chambers (Thermo Fisher Scientific) ∼24 hours before transfection. Constructs were transfected using FuGene HD Transfection Reagent for ∼72 hrs. Prior to imaging, cell culture media was replaced with complete imaging media (DMEM without phenol red, Thermo Fisher Scientific #31053028). For three-color experiments, cells were transfected with FASTKD2-mScarlet and POLRMT-HaloTag. Prior to imaging, cells were labelled with 200 nM Janelia Fluor 646 HaloTag ligand (Promega, #GA1120) for ∼1 hour in imaging media, then rinsed twice with imaging media, and finally, mtDNA was labelled with 0.01% v/v PicoGreen for an additional hour prior to imaging.

### Microscopy

Super-resolution microscopy was performed using a Carl Zeiss LSM980 inverted confocal microscope (Axio Observer 7) equipped with an Airyscan 2.0 detector and controlled by ZEN software Zen Blue version 3.6.095.03000. For fixed samples, images were acquired using a Plan-Apochromat 63X/1.4 NA oil objective with 405, 488, 561 and 647 nm laser lines. *Z*-stacks were collected in frame-switch mode with a 0.13 μm z-slice and 4X scan zoom and processed using Airyscan Joint Deconvolution (10 iterations). For three-color live-cell imaging, image acquisition was performed in line-switch mode with laser lines 488, 561, and 647 nm at 6X scan zoom. Images were processed using standard Airyscan Processing. Slides with 200 nm fluorescent beads (TetraSpeck™ Microspheres, 0.2 μm, Thermo Fisher Scientific) were imaged to perform channel alignment (Zen Black 3.0 SR FP2) on all processed images

### Image Analysis

Fiji (version 2.14.0/1.54p) and Zen software (Zen Blue version 3.6.095.03000 and Zen Lite version 3.8.99.01000) were used for visualization and figure generation of all imaging data. Super-resolution processed images were analysed using the mt-condensate pipeline to identify punctum features, including subpixel *X* and *Y* centroids using python (version 3.12.7)^3^. Puncta were segmented with custom threshold intensities for each channel for each replicate. Puncta were further linked in *Z* to generate 3D representations. Subpixel *Z*-centroid position was determined for each droplet by applying a gaussian fit. Nearest neighbour analysis was performed utilizing OpenCV to generate kd-tree data structures for each channel^4^. Probability distributions of distances for each channel pairing were plotted on a semi-log plot. Distances above 5 microns were excluded in plots and further calculations of statistical parameters. Correlation coefficients were calculated to identify the likelihood of the unpaired channels having similar distances as the paired channels.

To determine the average intensity of POLRMT, images were analysed using the above mt-condensate pipeline to identify the 3D coordinates of each punctum. Centroid data based on the *Z*-slice with the highest intensity of a given punctum was used to extract intensity values in the remaining channels. Intensities for each channel were normalized between 0 and 1, and all intensity arrays corresponding to a channel pair were subsequently averaged and visualized as 2D heatmaps. A separate script was utilized to extract the central 1D intensities from averaged 2D arrays for all replicates considering mtDNA and mtRNA as reference channels and plotted as mean intensities **±** S.D.

Line profiles for IMT1B treatment were performed using Fiji. Lines (∼0.6 μm) were drawn for selected ROI in representative images for both control and treatment group. Intensities were extracted using the ‘Plot Profile’ function with Visualization_toolset (Multichannel plot tool). A custom python script was used to calculate and plot normalized intensities per channel for both groups.

For 2D tracking, live imaging movies were first cropped to visualize events with isolated puncta in both channels for mt-nucleoids and MRGs. Cropped time-lapse images were next processed with a Gaussian filter (sigma = 1.6 in *X* and *Y*) to smooth out the puncta using Zen software. Lastly, puncta were tracked in 2D with trackpy (version 0.6.4) in Python^4^. *XY* tracks based on centroid position of mt-nucleoids (mtDNA, PicoGreen) and MRGs (FASTKD2-mScarlet) were used to plot 2D Euclidean distances between pairs of condensates over time.

## Supplemental Video Legends

**Supplemental Video 1**: Airyscan *Z*-stack of a fixed HeLa cell for an identified pair of mixed mt-nucleoids (yellow, anti-DNA) and MRGs (cyan, anti-GRSF1) relative to steady state ND6 mtRNA (magenta, RNA FISH). Scale bar = 200 nm.

**Supplemental Video 2**: Airyscan time-lapse of dynamic mixing, wetting and demixing of mt-nucleoids (yellow, Picogreen), MRGs (light blue, FASTKD2-mScarlet) and POLRMT (magenta, POLRMT-HaloTag- Janelia Fluor 646 ligand) in a live HeLa cell. Scale bar = 500 nm.

---

**Extended Fig. 1:**
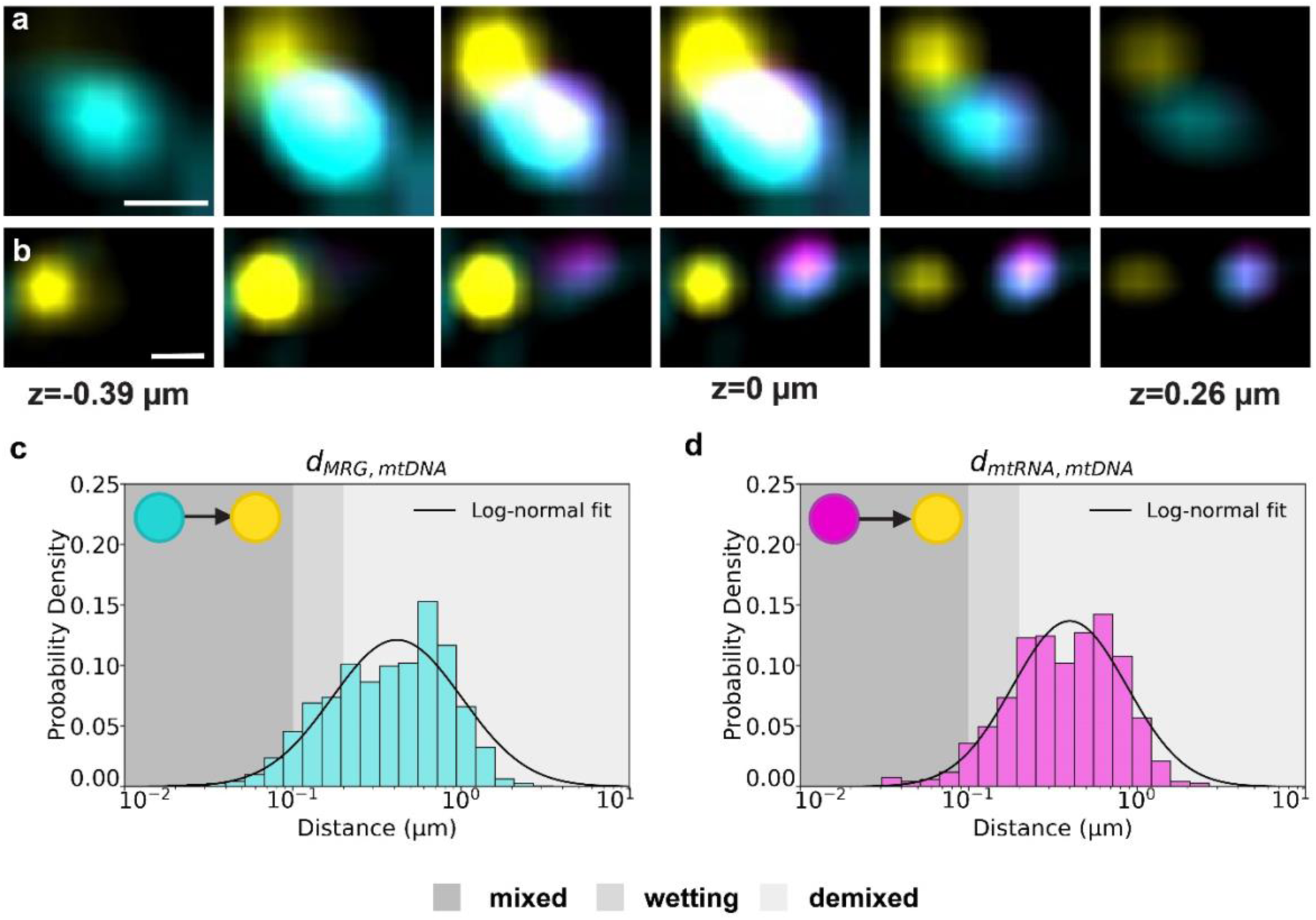
Endogenous localization of mitochondrial condensates and steady-state mtRNA. **a,b**, *Z*-stack of the wetting condensates (Fig. 1c) and demixed condensates (Fig. 1d), respectively, spanning ∼0.65 µm: mt-nucleoids (anti-DNA, yellow), MRGs (anti-GRSF1, cyan), and ND6 mtRNA (RNA FISH, magenta). Scale bar = 200 nm. **c-d,** Quantitative analysis of nearest neighbor distances between pairs of mitochondrial components. Semi-log plots of probability density distributions of nearest-neighbor distances: nearest mtDNA punctum to MRG **(c)** and nearest mtDNA punctum to ND6 mtRNA **(d)**. Shading of dark gray, medium gray and light gray indicate mixed, wetting and demixed populations, respectively, of the pairs of mitochondrial components within each plot (n=8 cells).

**Extended Fig. 2:**
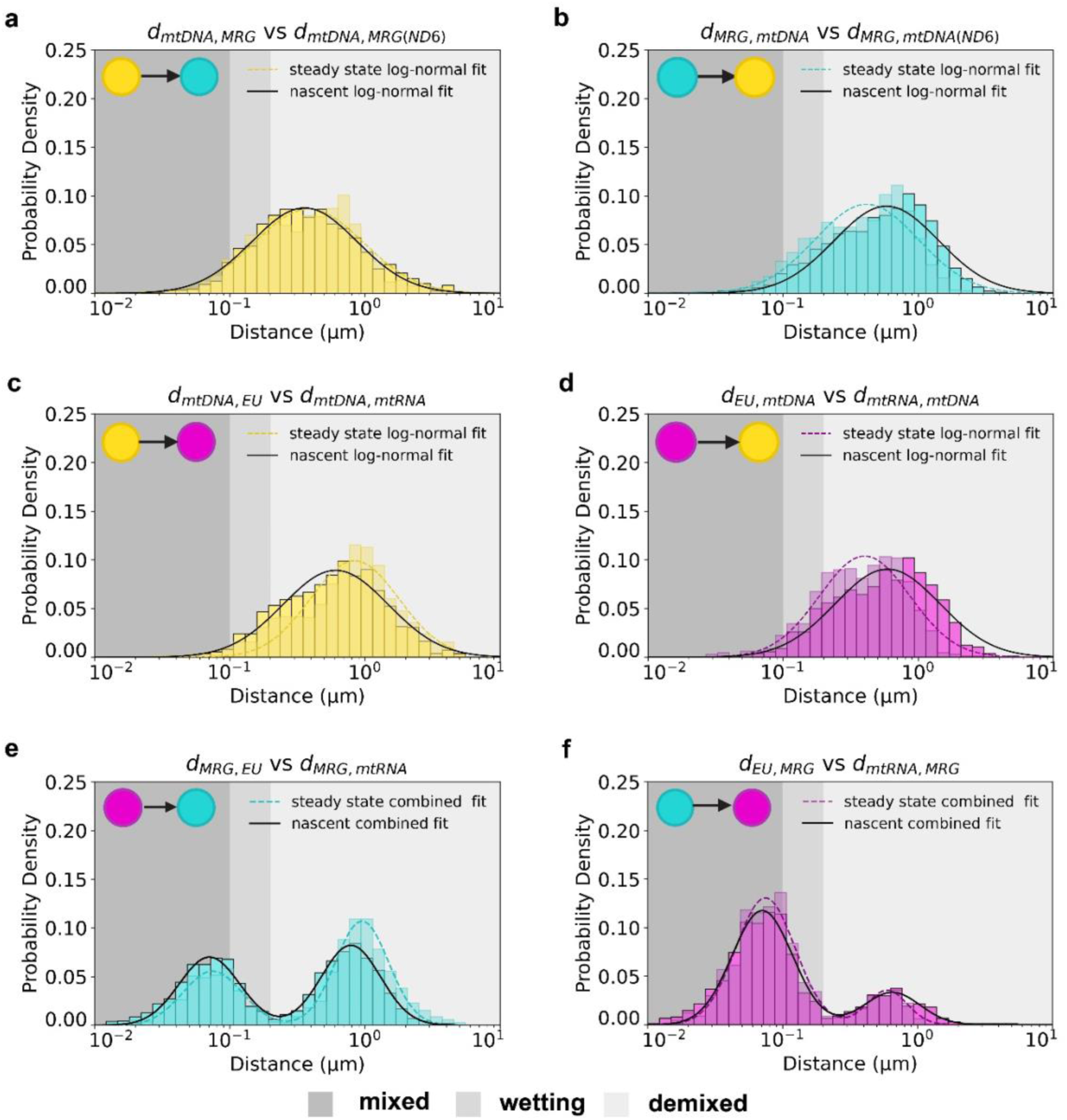
Comparison of nearest neighbor distances for EU- and ND6-labelled transcripts. **a-f**, Quantitative analysis of nearest neighbor distances between pairs of mitochondrial components for EU-labelled (nascent) and ND6-labelled (steady-sate) transcripts. Semi-log plots of probability density distributions of nearest-neighbor distances of EU-labelled mtRNA data (opaque, nascent) and ND6-labelled mtRNA (transparent, steady-state): mtDNA puncta to nearest MRG **(a);** MRG puncta to nearest mtDNA **(b);** mtDNA puncta to nearest mtRNA **(c)**; mtRNA puncta to nearest mtDNA **(d);** MRG puncta to nearest mtRNA**(e);** and mtRNA puncta to nearest MRG **(f)**. Shading of dark gray, medium gray and light gray indicate mixed, wetting and distal populations, respectively, of the pairs of mitochondrial components within each plot. For EU data, n=10 cells, for ND6 data, n=8 cells. P-values for nearest neighbor distance comparisons between nascent (EU) and steady-state (ND6) distributions are as follows: (**a**) p=0.39 for n=1,111 mtDNA puncta (EU), n=1,487 mtDNA puncta (ND6) (**b**) p=9.3E-29 for n=1,737 MRG puncta (EU), n=1,576 MRG puncta (ND6); (**c**) p=6.8E-10 for n=1,112 mtDNA puncta (EU), n=1,466 mtDNA puncta (ND6); (**d**) p=6.0E-24 for n=1,037 mtRNA puncta (EU), n=668 mtRNA puncta (ND6). Panels (**a-d**) were analyzed using two-tailed Welch’s t-test. (**e**) p=5.1E-26 for n=1,739 MRG puncta (EU), n=1,568 MRG puncta (ND6) and (**f**) p=0.94 for n=1035 mtRNA puncta (EU), n=668 mtRNA puncta (ND6) were analyzed using two-tailed Mann-Whitney U test.

**Extended Fig. 3:**
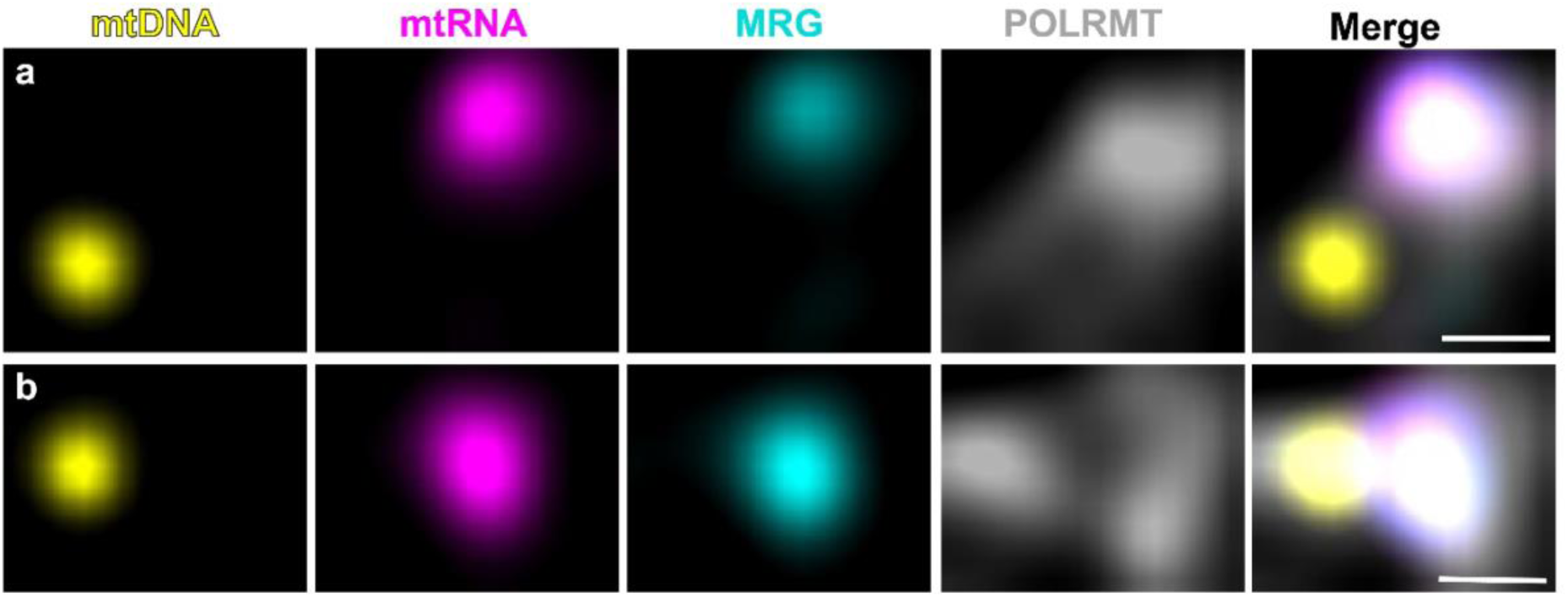
Colocalization of co-existing mitochondrial condensates and POLRMT. **a,b**, Single *Z*-slice representative AiryScan images of mt-condensates in which POLRMT is co-localizing exclusively with MRG and absent from an mt-nucleoid **(a)** and POLRMT is co-localizing with a pair of de-mixed mt-nucleoids and MRGs **(b)** in fixed wildtype HeLa cells over-expressing POLRMT-HaloTag (Oregon-Green-ligand, gray) with labelled ND6 mtRNA (RNA FISH, magenta), mt-nucleoids (anti-DNA, yellow), and MRGs (anti-GRSF1, cyan). Single channel of mtDNA, mtRNA, MRG, and POLRMT followed by a merged image. Scale bar = 200 nm. Intensities are individually adjusted for both panels for visualization (n=10 cells).

**Extended Fig 4:**
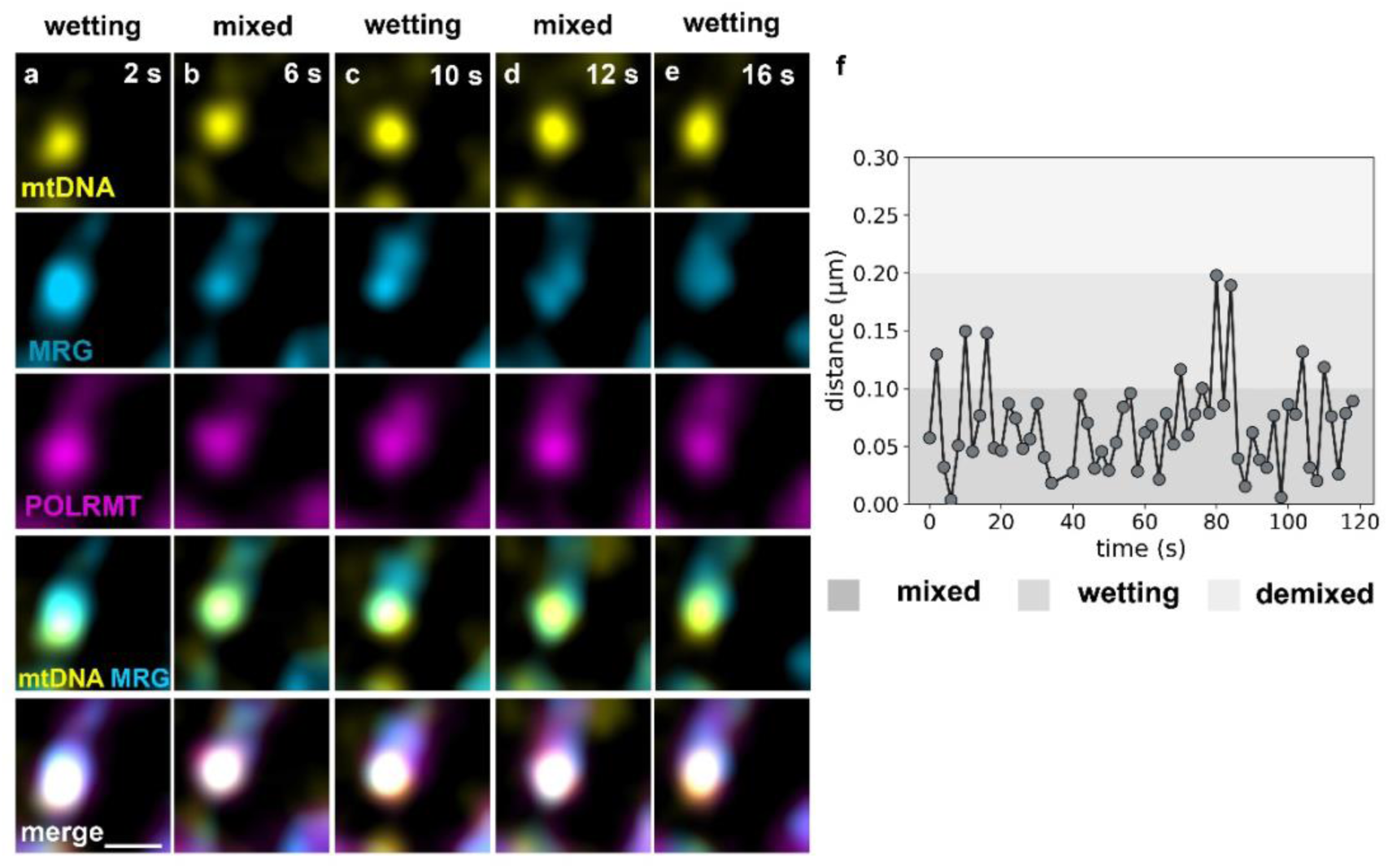
Dynamic organization of mitochondrial condensates in live cells. **a-e** Single *Z*-slice, Airyscan time-lapse imaging of live wildtype HeLa cell with three labelled components—mt-nucleoids (PicoGreen, yellow), MRGs (FASTKD2-mScarlet, light blue), and mtRNA polymerase (POLRMT-HaloTag, magenta) dynamically sampling mixing and wetting behaviors. Scale bar = 500 nm. **f,** Quantification of change in 2D Euclidean distance between centroids with time of a representative pair of mt-condensates (mt-nucleoid and MRG) from **a-e** (n=14 cells).

**Extended Fig. 5:**
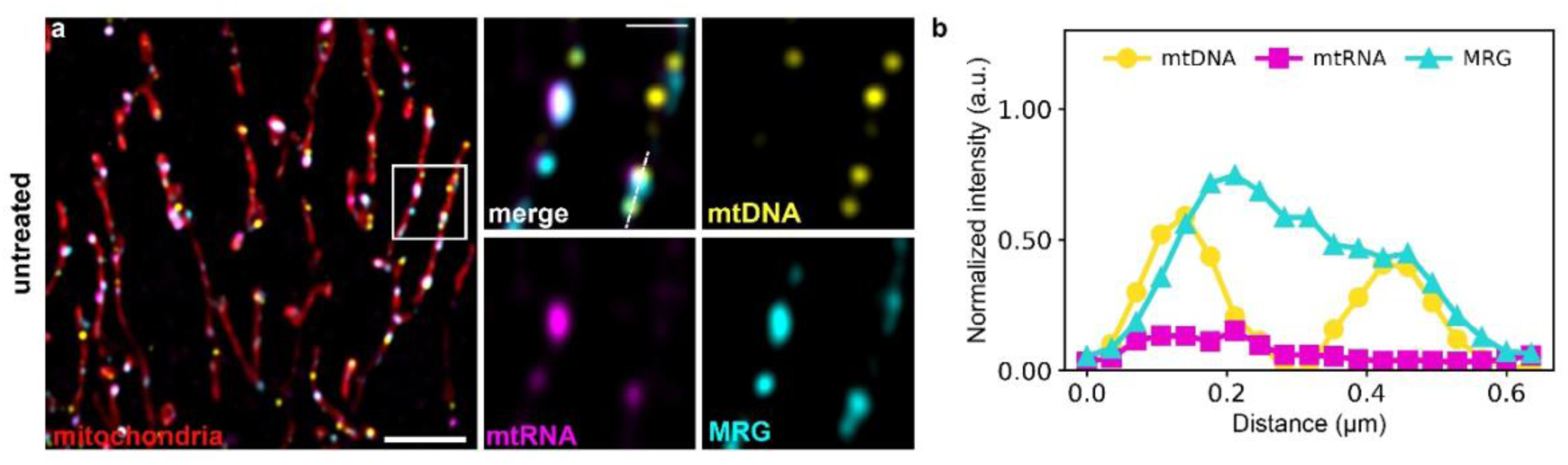
Control for dissolution of MRG with IMT1B. **a**, Single *Z*-slice of a representative Airyscan image of an untreated, fixed wildtype HeLa cell (negative control) with labelled mitochondrial network (MitoTracker Red, red), mt-nucleoids (anti-DNA, yellow), ND6 mtRNA (RNA FISH, magenta) and MRGs (anti-GRSF1, cyan). Scale bar = 2 μm. White box indicate regions of interest. Zoomed in image includes a three-channel overlay and single channels of mtDNA, mtRNA and MRGs. Scale bar = 500 nm. **b,** Line profile indicates normalized intensity distributions of mtDNA, mtRNA, and MRGs corresponding to dashed line on zoomed in ROI from **a**. Normalization with respect to DMSO control and IMT1B images (Fig. 5) (n=10 cells).

